# Phosphorylation Status Of MUS81 Is A Modifier Of Olaparib Sensitivity In BRCA2-Deficient Cells

**DOI:** 10.1101/2022.08.13.503764

**Authors:** Francesca Blandino, Eva Malacaria, Carolina Figlioli, Alessandro Noto, Giusj Monia Pugliese, Annapaola Franchitto, Pietro Pichierri

**Author notes:** Authors to whom correspondence should be addressed: Pietro Pichierri. Equally-contributing authors (Carolina Figlioli, Alessandro Noto, Giusj Monia Pugliese).

## Abstract

The MUS81 complex is crucial for preserving genome stability through resolution of branched DNA intermediates in mitosis and also for the processing of deprotected replication forks in BRCA2-deficient cells. Because of the existence of two different MUS81 complexes in mammalian cells that act in M or S-phase, whether and how the PARPi sensitivity of BRCA2-deficient cells is affected by loss of MUS81 function is unclear.

Here, using a mutant of MUS81 that impairs its function in M-phase, we show that viability of BRCA2-deficient cells but not their PARPi sensitivity requires a fully-functional MUS81 complex in mitosis. In contrast, expression of a constitutively-active MUS81 is sufficient to confer PARPi resistance. From a mechanistic point of view, our data indicates that deregulated action of the mitotic active form of MUS81 in S-phase leads to the cleavage of stalled replication forks before their reversal, bypassing fork deprotection, and engaging a Polθ-dependent DSBs repair.

Collectively, our findings describe a novel mechanism leading to PARPi resistance that involves unscheduled MUS81-dependent cleavage of intact, unreversed replication forks. Since this cleavage occurs mimicking the phosphorylated status of S87 of MUS81, our data suggest that hyperphosphorylation of this residue in S-phase might represent a novel biomarker to identify resistance to PARPi.

## INTRODUCTION

Mutations in BRCA1, BRCA2 and other BRCA genes characterize hereditary breast and ovarian cancer (1, 2). However, mutations in BRCA and homologous-recombination-related genes have been found also in subsets of sporadic breast cancers and, recently, also in prostate cancers (2, 3).

Mutations in BRCA genes confer elevated genome instability but also sensitize cancer cells to cisplatin and, more importantly, to PARP inhibitors, such as Olaparib (1, 4).

Mechanistically, sensitivity to PARPi would correlate with inability to properly repair or tolerate DSBs generated from the replication through single-strand lesions that are usually handled by PARP1-dependent pathways or trapped PARP/PARPi complexes (5, 6). In addition, sensitivity to PARPi in BRCA-defective backgrounds has been recently correlated with inability to protect stalled replication forks from degradation and to prevent accumulation of template gaps (7, 8).

Although PARPi proved to be very effective targeted therapeutic approaches, multiple mechanisms of resistance have been reported (6, 9). Given the relevance of targeted therapy of cancer, a detailed understanding of the possible routes to chemoresistance is fundamental for an educated therapeutic selection.

Up to now, the best characterised mechanism of resistance is the restoration of homologous recombination by either reconstitution of a functional BRCA1/2 allele or mutations that inactivate barriers to end-protection (6). Most recently, an additional route to PARPi resistance have been identified and involves the restoration of fork stability at reversed replication forks (10). In lower and higher eukaryotes, the Mus81/Mms4^EME1^ heterodimer, the MUS81 complex, is the major endonuclease involved in the resolution of recombination or replication DNA intermediates (11–13). The main physiological function of the MUS81 complex is performed during the G2/M phase of the cell cycle (14). However, a lot of evidence shows activation of the MUS81 complex also in S-phase under conditions of persisting replication stress or at deprotected forks (15–19). Interestingly, loss of MUS81 function at deprotected forks reduces chemosensitivity of BRCA2-deficient cells (20). However, loss of MUS81 is synthetic sick in a BRCA2-deficient mammary epithelial cells and seems to act synergistically with cisplatin and PARPi, increasing cell death (21, 22).

Such apparent paradox might be explained with the presence of two distinct MUS81 complexes in human cells, MUS81/EME1 and MUS81/EME2 (23). Thus, a detailed investigation of which of those two complexes is responsible for the synthetic-sick relationship with BRCA2 deficiency or with sensitivity to PARPi would require specific separation-of-function alleles of MUS81. We recently reported that CK2-dependent phosphorylation of MUS81 at S87 stimulates association with SLX4 in late G2/M and regulates the function of the MUS81/EME1 complex in mitosis (24). Abrogation of phosphorylation through expression of an unphosphorylable S87A-MUS81 allele impairs only mitotic resolution of DNA intermediates, making this form of MUS81 a separation-of-function mutant (24).

Here, we found that mitotic function of MUS81 is responsible for the observed synthetic sick MUS81/BRCA2 interaction, but is not involved in the Olaparib resistance associated with BRCA2 loss. Most importantly, we found that deregulated function of MUS81/EME1 complex in S-phase is sufficient to revert Olaparib sensitivity in BRCA2-deficient backgrounds through the cleavage of stalled forks before fork reversal and degradation, and subsequent involvement of a Polθ-dependent pathway of DSBs repair. Therefore, our results represent the first demonstration of regulatory phosphorylation of the MUS81 subunit as a novel mechanism of chemoresistance to PARPi in BRCA2-deficient cells.

## MATERIALS AND METHODS

### Cell Cultures

The SV40-transformed MRC5 fibroblast cell line (MRC5SV40 or simply MRC5 in the text) was a generous gift from Dr. P. Kannouche (IGR, Villejuif, France). The cells were maintained in Dulbecco’s modified Eagle’s medium (DMEM; Life Technologies) supplemented with 10% FBS (Boehringer Mannheim) and incubated at 37 °C in a humidified 5% CO2 atmosphere. Cell line were routinely tested for mycoplasma contamination and maintained in cultures for no more than one month.

MRC5shMUS81 cells and generation of stable clones complemented with MUS81 mutants were described in Palma et al (24).

To obtain MUS81 knockout in MFC7 (breast cancer) and MRC5 cell lines, two RNP complexes obtained with Alt-R CRISPR-Cas system (IDT) specific to exon 1 of MUS81 were nucleofected using the Neon Trasfection System. For targeting of MUS81 gene the oligos were: 5’TCCTACAGCACTTCGGAGAC3’ and 5’CATGCCCCGGACTCACCATC3’. Clones were isolated, verified by Western blotting and then by Sanger sequencing. Nuclease-inactivating mutations D338A/D339A were introduced in the S87D-MUS81 sequence by site-directed mutagenesis in the pIRES-neo-Flag-MUS81 plasmid (24). After confirmation, the expressing plasmid was transfected into MRC5 MUS81 KO cells to generate stable cell lines.

### Chemicals

Hydroxyurea (Sigma-Aldrich) was used at 2 mM; MIRIN, the inhibitor of MRE11 exonuclease activity (Calbiochem), was used at 45 µM. Cells were accumulated in mitosis upon Nocodazole (Sigma-Aldrich) treatment for 24 hours at 0.5 μg/μl. To label parental ssDNA, 5’-Iodo-2’deoxyuridine (IdU-Sigma-Aldrich) was used for 20 hours at 100μM. To label DNA fibers: 5’-Chloro-2’-deoxyuridine (CldU-Sigma-Aldrich) was used at 200μM, IdU was used at 50μM for 20 minutes.

### Plasmids and RNA interference

The RuvA-GFP-NLS expression plasmid has been described in Malacaria et al (25). RNA interference against BRCA2 was performed using ON-TARGETplus Human BRCA2-SMARTpool (Dharmacon). For MUS81 interference it was used FlexiTube HsMUS81 6 (Qiagen). SMARCAL1 was depleted using the MISSION esiRNA HUMAN SMARCAL1 (Sigma-Aldrich). In all experiments, cells were transfected using Interferin (Polyplus). The efficiency of protein depletion was monitored by Western blotting 48-72h after transfection.

### Viability Assays

Clonogenic assay was performed seeding 1000 cells in 60-mm Petri dish. Colonies were fixed in methanol-acetic acid 3:1 and stained with 5mg/ml GIEMSA (Sigma). Cell survival was expressed relative to control cells. For MTT assay, 24h after RNA interference, cells were plated in 96-well plate; at the end of treatment, viability was quantified as manufacture protocol (MTT, Abcam, ab211091).

### Immunofluorescence

Immunofluorescence microscopy was performed on cells grown on 35-mm cover-slips and harvested at the indicated times after treatments. For detection of parental ssDNA, cells were labeled with 50µM Iododeoxyuridine (IdU, Sigma-Aldrich) for 36 hours, then release for 2 h in fresh culture media. After treatments, cells were washed with PBS and permeabilized with 0.5% Triton-X 100 and fixed with 3% PFA-2% sucrose in PBS on ice for 10 min. After blocking in 3% BSA/PBS for 15 min, staining was performed with anti-IdU antibody (mouse-monoclonal anti-BrdU/IdU; clone b44 Becton Dickinson, 1:80) for 1h at 37°C in a humidified chamber. Alexa Fluor® 488 conjugated-goat anti-mouse were used at 1:200. Nuclei were stained with 4’,6-diamidino-2-phenylindole (DAPI). Detection of ɣ-Tubulin or pS87 MUS81 was performed using anti-ɣ-Tubulin (1:200, Sigma-Aldrich) or anti-pS87MUS81 (WB 1:1000, IF 1:200, Abgent (24)).

Coverslips were observed at 40× objective with the Eclipse 80i Nikon Fluorescence Microscope, equipped with a VideoConfocal (ViCo) system.

For detection of SMARCAL1 and RAD51, immunofluorescences were performed as above (anti-SMARCAL1, Abcam, 1:100; anti-RAD51, Abcam, 1:1000) and nuclei were stained with DAPI.

For immunodetection of BRCA2 foci, cells were pre-extracted with CSK buffer on ice, then fixed in 2%PFA for 15min. After blocking in 10% FBS/PBS, cells were incubated with anti-BRCA2 antibody (rabbit-polyclonal, Bethyl, 1:200) for 1h at 37°C in a humidified chamber. Alexa Fluor® 594 conjugated-goat anti-rabbit were used at 1:200. Nuclei were stained with DAPI and foci were acquired randomly using 60x objective.

For each time point at least 200 nuclei were examined. Parallel samples incubated with either the appropriate normal serum or only with the secondary antibody confirmed that the observed fluorescence pattern was not attributable to artefacts.

For experiments with labeling cellular DNA with EdU (5-ethynyl-2’-deoxyuridine), EdU was added to the culture media (10μM), for 30 min. Detection of EdU was performed used Click-it EdU imaging Kits (Invitrogen).

### DNA Fiber Analysis

Cells were pulse-labelled with 50 µM 5-chloro-2’-deoxyuridine (CldU) and 250 µM 5-iodo-2’-deoxyuridine (IdU) as indicated, with or without treatment as reported in the experimental schemes. DNA fibers were spread out as previously reported (26). For immunodetection of labelled tracks the following primary antibodies were used: anti-CldU (1:60; rat-monoclonal anti-BrdU/CldU; BU1/75 ICR1 Abcam) and anti-IdU (1:10; mouse-monoclonal anti-BrdU/IdU; clone b44 Becton Dickinson). The secondary antibodies were goat anti-mouse Alexa Fluor 488 or goat anti-rat Alexa Fluor594 (Invitrogen). The incubation with antibodies was accomplished in a humidified chamber for 1 h at 37°C. Images were acquired randomly from fields with untangled fibers using Eclipse 80i Nikon Fluorescence Microscope, equipped with a Video Confocal (ViCo) system. The length of labelled tracks was measured using the Image-Pro-Plus 6.0 software. A minimum of 100 individual fibers were analyzed for each experiment and the mean of at least three independent experiments presented. Statistics were calculated using GraphPad Prism Software.

### In situ PLA for ssDNA–protein interaction

The *in situ* proximity-ligation assay (PLA, Duolink kit from Sigma-Aldrich) was performed according to Iannascoli et al (26). Briefly, for parental ssDNA, cells were labeled with 100µM IdU for 24 hours before treatments. Antibody staining was carried out according to the standard immunofluorescence procedure. The primary antibodies used were: rabbit-polyclonal anti-RAD51 (Abcam), mouse-monoclonal anti-BrdU/IdU (clone b44 Becton Dickinson).

The negative control consisted of using only one primary antibody. The incubation with all antibodies was accomplished in a moist chamber for 1 h at 37 °C. Samples were incubated with secondary antibodies conjugated with PLA probes MINUS and PLUS: the PLA probe anti-mouse PLUS and anti-rabbit MINUS. Samples were mounted in ProLong Gold antifade reagent with DAPI (blue). Images were acquired randomly using Eclipse 80i Nikon Fluorescence Microscope, equipped with a VideoConfocal (ViCo) system, at least 200 nuclei were examined and foci were scored at 40×.

### Quantitative in situ analysis of protein interactions at DNA replication forks (SIRF)

Cells were incubated with 125µM EdU for 8 minutes before treatments with HU. After treatment, cells were fixed with 2%PFA/PBS for 15 min at room temperature (RT). Cells were next permeabilized with 0.25%Triton-X100 in PBS for 15 minutes at RT. Detection of EdU was performed used Click-it EdU imaging Kits as described in (27). After Click-It reaction, slides were incubated with the primary antibodies: anti-biotin (mouse, Invitrogen, 1:200) and anti-SMARCAL1 (rabbit, Abcam, 1:200).

### Neutral Comet Assay

DNA breakage induction was evaluated by Comet assay (single cell gel electrophoresis) in non-denaturing conditions as described in Murfuni et al (28). Briefly, dust-free frosted-end microscope slides were kept in methanol overnight to remove fatty residues. Slides were then dipped into molten agarose at 1% and left to dry. Cell pellets were resuspended in PBS and kept on ice to inhibit DNA repair. Cell suspensions were rapidly mixed with LMP agarose at 0.5% kept at 37 °C and an aliquot was pipetted onto agarose-covered surface of the slide. Agarose embedded cells were lysed by submerging slides in lysis solution (30 mM EDTA, 0,5% SDS) and incubated at 4 °C, 1 h in the dark. After lysis, slides were washed in TBE 1X running buffer (Tris 900 mM; boric acid 900 mM; EDTA 20 mM) for 1 min. Electrophoresis was performed for 18 min in TBE 1X buffer at 20 V/6-7mA. Slides were subsequently washed in distilled H2O and finally dehydrated in ice cold methanol. In some cases, Comet assay was performed on cells pulse-labelled with 50µ IdU 30min before treatment to analyse DSBs in S-phase cells. In this case, slides were denatured under acidic conditions before undergoing detection of IdU using anti BrdU antibody. Nuclei were stained with GelRed (1:1000) and visualized with a fluorescence microscope (Zeiss), using a 40X objective, connected to a CCD camera for image acquisition. At least 300 comets per cell line were analyzed using CometAssay IV software (Perceptive instruments) and data from tail moments processed using Prism software. Apoptotic cells (smaller Comet head and extremely larger Comet tail) were excluded from the analysis to avoid artificial enhancement of the tail moment. A minimum of 200 cells was analyzed for each experimental point.

### Western Blotting Analysis

Western blots were performed using standard methods. Blots were incubated with primary antibodies commercially obtained against: anti-MUS81 (1:1000; Santa Cruz Biotechnologies), anti-SMARCAL1 (1:2000; Abcam), anti-RAD51 (1:5000; Abcam), anti-BrdU (1:50, Becton Dickinson), anti-BRCA2 (1:1000; Bethyl), anti-Lamin B1 (1:10000; Abcam), GFP (1:1000; Santa Cruz Biotechnology), GAPDH (1:5000; Millipore). HRP-conjugated matched secondary antibodies were from Jackson Immunoresearch and were used at 1:40000.

Horseradish peroxidase-conjugated goat specie-specific secondary antibodies (Santa Cruz Biotechnology, Inc.) were used. Quantification was performed on scanned images of blots using ImageJ software, and values are shown on the graphs as indicated.

### Chromatin Fractionation

Cells (4 × 10^6^ cells/ml) were resuspended in buffer A (10 mM HEPES, [pH 7.9], 10 mM KCl, 1.5 mM MgCl2, 0.34 M sucrose, 10% glycerol, 1 mM DTT, 50 mM sodium fluoride, protease inhibitors [Roche]). Triton X-100 (0.1%) was added, and the cells were incubated for 5 min on ice. Nuclei were collected in pellet by low-speed centrifugation (4 min, 1,300 ×g, 4°C) and washed once in buffer A. Nuclei were then lysed in buffer B (3 mM EDTA, 0.2 mM EGTA, 1 mM DTT, protease inhibitors). Insoluble chromatin was collected by centrifugation (4 min, 1,700 × g, 4°C), washed once in buffer B, and centrifuged again under the same conditions. The final chromatin pellet was resuspended in 2X Laemmli buffer and sonicated in a Tekmar CV26 sonicator using a microtip at 50% amplitude. After samples were boiled for 10 min at 95°C and then subjected to Western blot as reported below.

### Statistical Analysis

All the data are from at least two independent sets of experiments. Statistical comparisons were performed as indicated in the figure legends. P < 0.5 was considered significant.

## RESULTS

### Mitotic phosphorylation of MUS81 regulates viability in the absence of BRCA2

Depletion of MUS81 reduces viability of BRCA2-deficient cells (21). However, it is not known which of the roles of MUS81, resolution of late recombination intermediates in mitosis or processing of degraded forks in S-phase, is specifically required. Abrogation of CK2-dependent phosphorylation of MUS81 at S87 disrupts only the mitotic function of MUS81 (24). Hence, we decided to use the S87A MUS81 separation of function mutant to verify whether impaired resolution of DNA intermediates in mitosis was sufficient to affect viability of BRCA2-deficient cells.

We first analysed clonogenic survival following transient depletion of BRCA2 in stable shMUS81 cell lines complemented with wild-type MUS81 or the two S87 mutants that abrogates or mimics phosphorylation by CK2 (24). Transfection with BRCA2 siRNA led to comparable protein knockdown in all cell lines (Figure 1A) and reduced clonogenic survival when combined with loss of MUS81 (Figure 1B), in agreement with previous studies (21). Interestingly, the expression of the mitotic inactive, CK2-unphosphorylable, MUS81^S87A^ mutant was sufficient to recapitulate the effect of MUS81 depletion on the viability of BRCA2-deficient cells while survival was not affected by expression of the phosphomimetic MUS81^S87D^ (Figure 1B).

**Figure 1.**
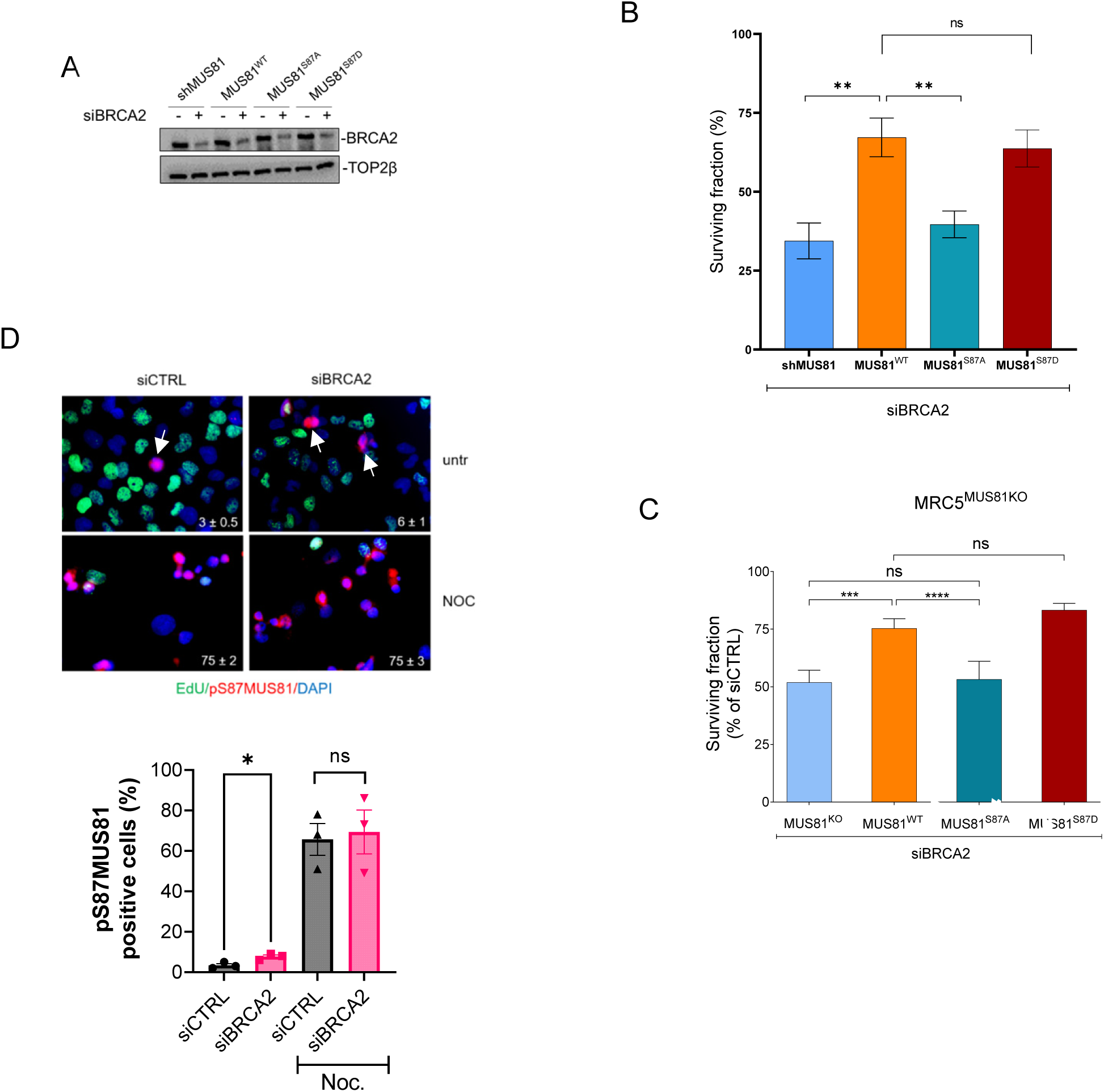
The mitotic function of MUS81 is sufficient to promote viability in the absence of BRCA2. (A) Western blotting analysis of BRCA2 protein depletion in shMUS81 MRC5 cells complemented with the indicated form of MUS81 48h after transfection with control or BRCA2 siRNAs. (B) shMUS81 MRC5 cells complemented with the indicated MUS81 form, or nothing (shMUS81) were depleted of BRCA2 and evaluated for clonogenic survival. The graph shows the percentage of surviving fraction normalized against the corresponding siCtrl-transfected cells. Data are presented as mean from at least quintuplicate experiments ± S.E. The image show a representative plate from the triplicate set. (C) Viability of MUS81 KO MRC5 cells transiently transfected with the indicated constructs and a siBRCA2 oligo was analysed by clonogenic survival. The graph shows the percentage of surviving fraction normalized against the corresponding siCtrl-transfected cells. (D) MUS81 phosphorylation evaluated by anti-pS87MUS81 immunofluorescence (red) after transfection with siCTRL or siBRCA2 oligos as in (A). S-phase cells were labelled by a pulse with EdU (green). Nuclei were stained with DAPI (blue). Cells were either left untreated (untr) or accumulated in M-phase with nocodazole (NOC). Numbers represent the mean value of pS87MUS81-positive cells ± SE. Arrows indicates pS87MUS81^+^ cells in asynchronous cultures. The graph shows quantification from three independent experiments (Student’s t-test)

To further substantiate our observation, we used Crispr to knock out MUS81 in MRC5SV40 cells (MRC5^KO^; Supplementary Figure 1A). These cells have been transiently complemented with wild-type MUS81 or its S87 phosphorylation mutants (Supplementary Figure 1B) and viability assessed by clonogenic assay upon depletion of BRCA2 (Figure 1C). Consistent with observation in shMUS81 cells, upon BRCA2 depletion, loss of S87 phosphorylation reduced viability just as the absence of MUS81. Compared to the wild-type MUS81, expression of the constitutively-active S87D MUS81 mutant in the MRC5^KO^ cells did not have any effect on survival of BRCA2-depleted cells.

These results suggest that S87-MUS81 phosphorylation could be elevated in BRCA2-depleted cells. To test this possibility, we performed immunofluorescence on asynchronous or mitotically-enriched populations of cells transiently depleted of BRCA2. Staining with the anti-pS87-MUS81 antibody (24) showed that the number of positive cells was doubled upon BRCA2 depletion in asynchronous cells (Figure 1D) and that S87-phosphorylated MUS81 did not co-localise with S-phase cells, as expected (24). In contrast, the percentage of pS87-MUS81-positive cells was similar in mitotically-enriched cells independently of the presence of BRCA2. This result suggests that loss of BRCA2 stimulates mitotic phosphorylation of MUS81 at S87 although this difference is not detectable when cells are enriched in mitosis and pS87 MUS81 phosphorylation is physiologically high (24).

Although the simple loss of mitotic MUS81 function in mitosis was sufficient to phenocopy the effect of the absence of the protein in BRCA2-deficient cells, it did not affect the presence of DSBs in cells depleted of BRCA2 (Supplementary Figure 2A, B). Furthermore, the loss of BRCA2 did not affect the elevated level of DSBs observed in the S87D-MUS81 mutant ((24); Supplementary Figure 2A, B). Consistent with previous data (24), the elevated formation of DSBs associated with the expression of S87D-MUS81 was detected in S-phase, either in the presence of BRCA2 or in its absence (Supplementary Figure 2C).

Altogether, results from the mitotic separation-of-function S87A MUS81 mutant indicates that is the mitotic function of MUS81 to be crucial for viability in BRCA2-depleted cells. Of note, unscheduled generation of DSBs at the fork caused by the presence of the deregulated MUS81-S87D mutant (24) does not synergize, in terms of viability, with loss of BRCA2.

### The status of mitotic phosphorylation of MUS81 affects chemosensitivity of BRCA2-deficient cells

Downregulation of MUS81 is associated to increased PARPi sensitivity (29). However, abrogating the MUS81-dependent cleavage at deprotected replication forks and restored fork protection reduces chemosensitivity in BRCA2-deficient cells (7, 20). Since MUS81 may perform multiple functions, we exploited the S87A separation-of-function MUS81 mutant to determine which of the functions of MUS81 affects sensitivity to PARPi.

We first evaluated viability of shMUS81 cells expressing the two regulatory mitotic MUS81 mutants in response to the prototypical PARPi Olaparib. These cells were transfected with BRCA2 siRNA and their sensitivity tested by clonogenic assay after 24h treatment with 10µM of Olaparib and seeding at low density for 10 days, a condition expected to greatly affect survival in the absence of BRCA2 (Figure 2A and B). As expected, viability of untreated BRCA2-deficient cells was similarly reduced by depleting MUS81 or abrogating its mitotic function by the S87A mutation. In response to Olaparib, the abrogation of the mitotic function of MUS81 or its depletion slightly reduced viability compared to the wild-type MUS81. However, the expression of the constitutively active S87D mitotic mutant of MUS81 apparently conferred Olaparib resistance to BRCA2-deficient cells.

**Figure 2.**
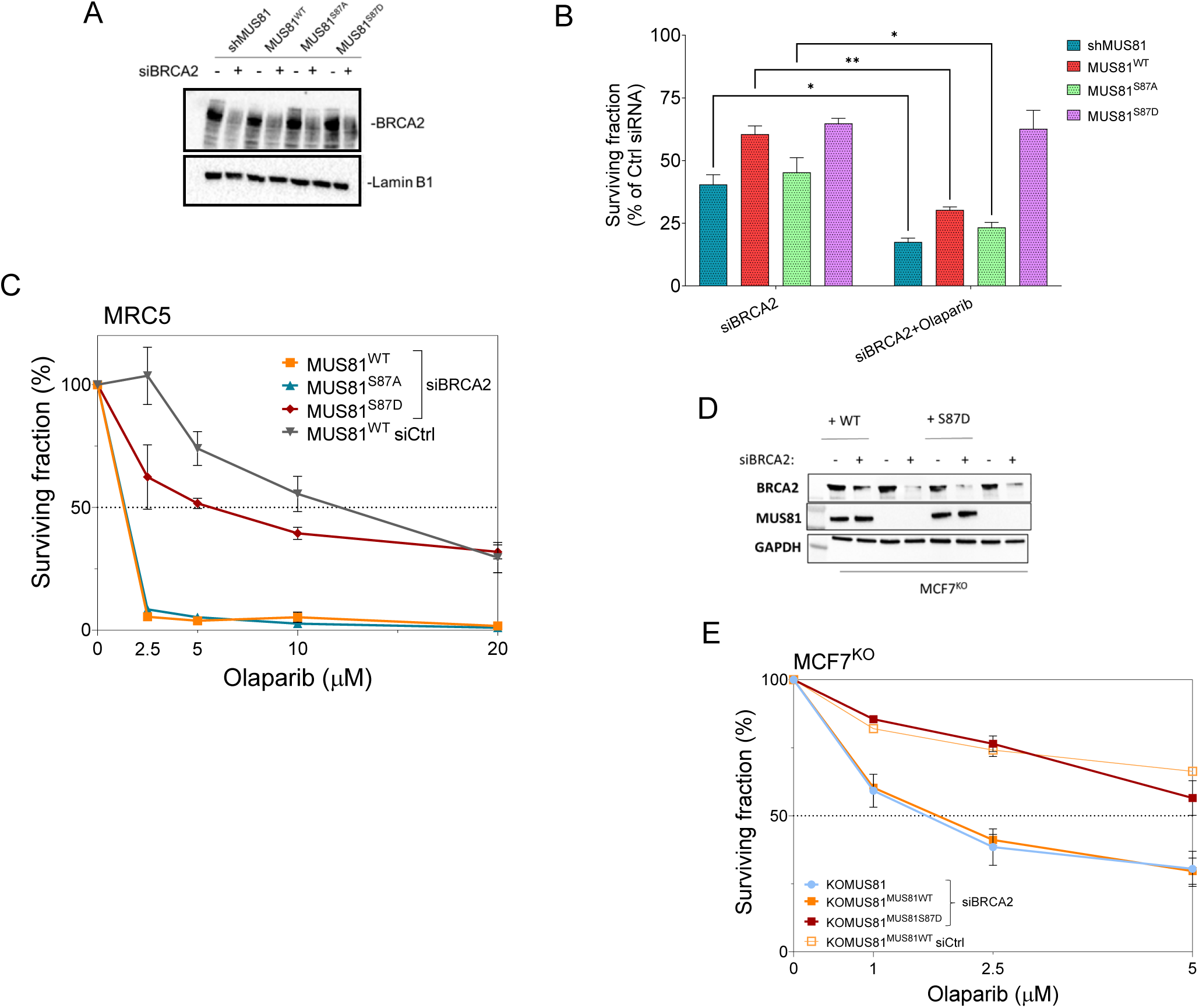
Constitutive phosphorylation of MUS81 at S87 confers resistance to Olaparib treatment in BRCA2-deficient cells. (A) Western blotting analysis of BRCA2 protein depletion in shMUS81 MRC5 cells complemented with the indicated form of MUS81 48h after transfection with control or BRCA2 siRNAs. (B) Cells were treated with Olaparib 24h after transfection with Ctrl or BRCA2 siRNAs and viability assessed by clonogenic survival. The graph shows the percentage of surviving fraction normalized against the respective siCtrl-transfected cells. Data are presented as mean from at least triplicate experiments ± S.E. (C). Short-term viability assay. Forty-eight hours after BRCA2 depletion, cells were seeded in 96-well plate and treated with increasing concentrations of Olaparib and tested by MTT assay. Data are presented as mean from at least triplicate experiments ± S.E. (D) Western blotting analysis of BRCA2 protein depletion in MUS81 KO MCF7 cells (MCF7^KO^) complemented with the indicated form of MUS81 48h after transfection with control or BRCA2 siRNAs. (E) Forty-eight hours after BRCA2 silencing, MUS81 KO MCF7 cells were complemented with MUS81WT or S87D and treated with increasing concentrations of Olaparib. Data are presented as mean ± S.E. (n=3). Statistical analysis was performed by Student’s t-test. ns: not significant; *P<0.5; **P<0.1; ***P<0.001.

The S87D MUS81 mutant leads to DSBs in S-phase. To confirm that unscheduled cleavage by MUS81 in S-phase reverted Olaparib sensitivity of BRCA2-deficient cells, we evaluated viability by MTT assay over a range of doses (Figure 2C). As expected, loss of BRCA2 sensitized cells expressing wild-type MUS81 to Olaparib and this phenotype was barely affected by the S87A-MUS81 mutant, which abrogates the MUS81 complex function in mitosis. Nevertheless, short-term viability data confirmed that expression of the S87D-MUS81 mutant confers PARPi resistance to BRCA2-deficient cells.

We next investigated if the expression of S87D-MUS81 could attenuate Olaparib sensitivity also in breast cancer cells. Thus, we transiently expressed wild-type or S87D-MUS81 in the MCF7^KO^ breast cancer cells described in Suppl. Figure 1, and analysed sensitivity to Olaparib upon depletion of BRCA2 by siRNA transfection (Figure 2D, E). Although, in our hands, MCF7 cells were more resistant to Olaparib than MRC5, depletion of BRCA2 sensitised also MCF7 cells to treatment and MUS81 knock-out did not affect sensitivity. However, and consistent with data in MRC5 cells, expression of the S87D-MUS81 mutant greatly reduced Olaparib sensitivity in BRCA2-depleted MCF7 breast cancer cells.

Collectively, these results indicate that the selective inactivation of the mitotic function of MUS81 does not affect sensitivity to Olaparib, while the deregulated function of MUS81 in S-phase, as conferred by mimicking constitutive S87 phosphorylation, is sufficient to attenuate Olaparib sensitivity in BRCA2-depleted cells.

### The S87D MUS81 mutant reverts fork degradation in BRCA2-deficient cells by targeting stalled replication forks independently of fork reversal

Restoration of fork protection or inability to cleave deprotected forks is sufficient to confer PARPi or cisplatin resistance in BRCA2-deficient cells (7, 20). Our data demonstrate that the S87D MUS81 mutation is sufficient to greatly reduce Olaparib sensitivity in BRCA2-deficient cells. Although deregulated mitotic function of MUS81 did not affect replication fork progression under unperturbed conditions (Supplementary Figure 2D), we performed DNA fibre assay to analyse whether nascent strand degradation, which is linked to fork deprotection in the absence of BRCA2, was affected by the S87D-MUS81 mutant (Figure 3A).

**Figure 3.**
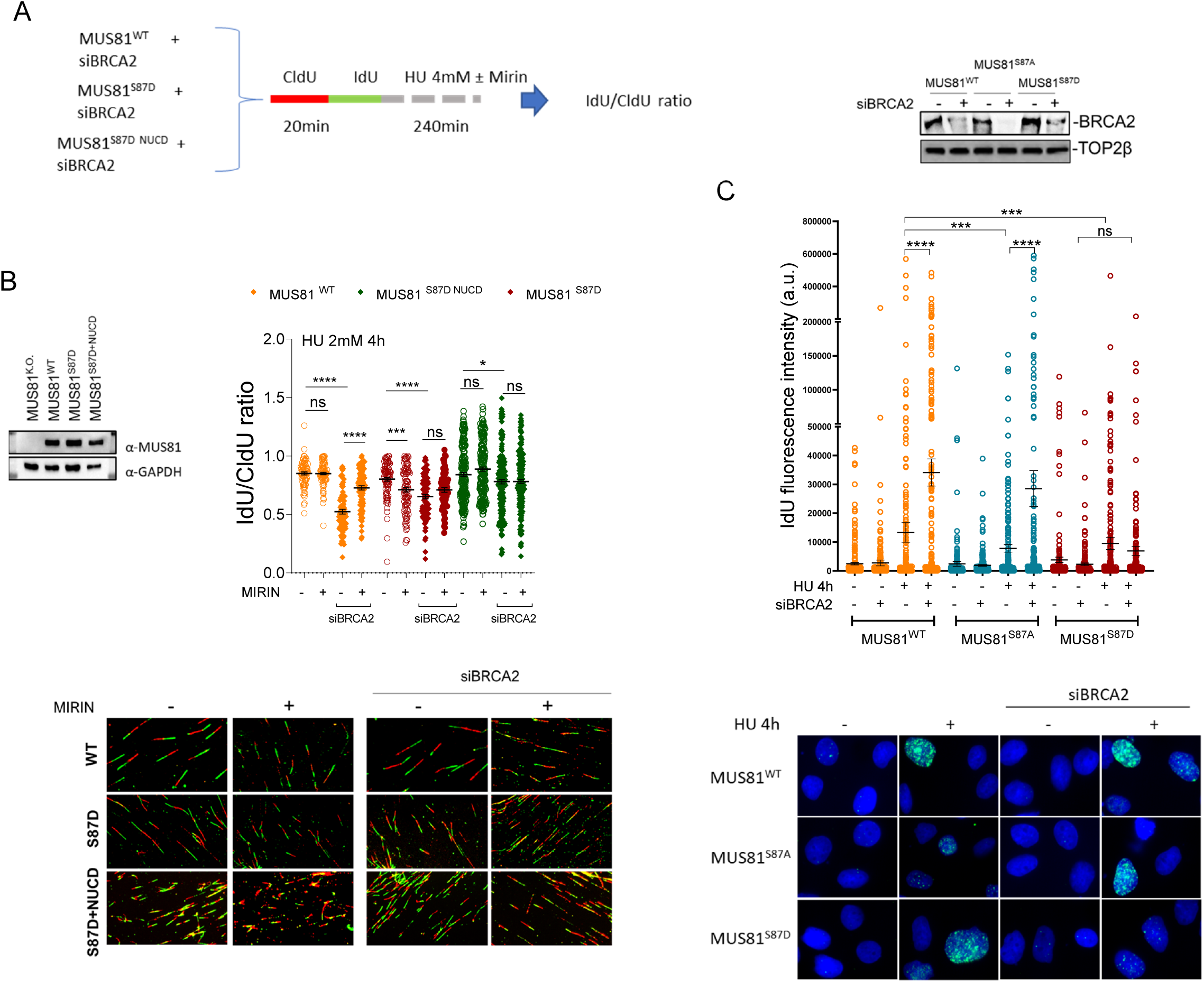
Constitutive phosphorylation of MUS81 at S87 prevents MRE11-dependent fork degradation in BRCA2-deficient cells. (A) Experimental scheme. (B) IdU/CldU ratio in BRCA2-depleted MRC5 KO MUS81 cells complemented with the indicated MUS81 mutant. S87D+NUCD = S87D mutant combined with the two nuclease-inactivating mutations (D338A/D339A); n ≥ 150 tracts were scored for each data set out of 3 repeats. Bars represent the mean ± S.E. WB shows the MUS81 expression in the (C) Unscheduled activity of MUS81 reduces the exposure of parental ssDNA early after fork arrest. Detection of ssDNA by native IF in MRC5 cells treated with 2mM HU. Graph shows the individual values from 3 independent replicates. The bars indicate mean ± SE. Representative images are shown. Statistical analyses were performed by Mann–Whitney test (ns, not significant; *P< 0.5; **P< 0.1; ***P< 0.01; ****P< 0.001).

In agreement with previous reports (30–32), depletion of BRCA2 induced fork deprotection in cells expressing wild-type MUS81. Indeed, BRCA2-depleted cells showed shorter IdU-labelled tracks after HU leading to a IdU/CldU ratio < 1 (Figure 3B). Inhibition of MRE11 with Mirin restored normal track length confirming the degradation phenotype (Fig. 3B). Compared with the wild-type, the S87D MUS81 mutant did not affect the IdU/CldU ratio, which was reduced by Mirin (Figure 3B). This unexpected IdU/CldU ratio reduction was rescued by abrogating nuclease activity (S87D+NCD), possibly indicating involvement of the deregulated function of MUS81 on replication substrates (Figure 3B). BRCA2-depleted cells expressing the S87D MUS81 mutant showed shorter IdU-labelled tracks as compared to control-depleted cells but inhibition of MRE11 did not rescue the phenotype suggesting that fork deprotection did not occur in the presence of the deregulated MUS81 mutant and this was not recovered by inactivating the nuclease activity (Figure 3B).

In the absence of BRCA2, degradation of the nascent strand at reversed forks results in exposure of parental ssDNA (33). Similarly, also the accumulation of parental gaps observed in BRCA2-deficient cells leads to persistent parental ssDNA (8). Since expression of S87D-MUS81 abrogated fork degradation, we next performed native IdU detection to analyse the presence of exposed parental ssDNA after HU (Figure 3C). In cells expressing wild-type MUS81, loss of BRCA2 resulted in elevated exposure of parental ssDNA, as expected. The presence of the mitotic-inactive MUS81 mutant decreased the amount of parental ssDNA in response to replication arrest, when BRCA2 is depleted. In contrast, while the expression of S87D MUS81 only induced minor changes in the amount of parental ssDNA exposed in CTRL RNAi-treated cells, it prevented its accumulation in response to HU in BRCA2-depleted cells, consistent with rescue of the fork degradation phenotype seen by DNA fibre assay.

In BRCA2-defective cells, fork deprotection and nascent-strand degradation occur before MUS81-dependent DNA breakage (31, 33, 34). Since the deregulated S87D MUS81 mutant introduced DSBs in S-phase irrespective of the presence of BRCA2 (Supplementary Figure 2B) but apparently prevented fork degradation in the absence of BRCA2 (Figure 3B), we investigated whether it could differentially affect the formation of DSBs in response to fork stalling.

Thus, we performed neutral Comet assay in cells transiently-depleted of BRCA2 (Figure 4A). As expected, 4h of HU did not result in a great increase of DSBs in cells expressing the wild-type MUS81 (15), while it raised the amount of DSBs of about 4-fold in BRCA2-depleted cells (Figure 4B). Of note, although the expression of the deregulated S87D MUS81 mutant greatly increased DSBs formation under unperturbed replication in control-depleted cells, it failed to further increase DSBs upon replication arrest, irrespective of BRCA2 depletion (Figure 4B), suggesting that unscheduled cleavage targeted all the potential DNA substrates already in the absence of HU.

**Figure 4.**
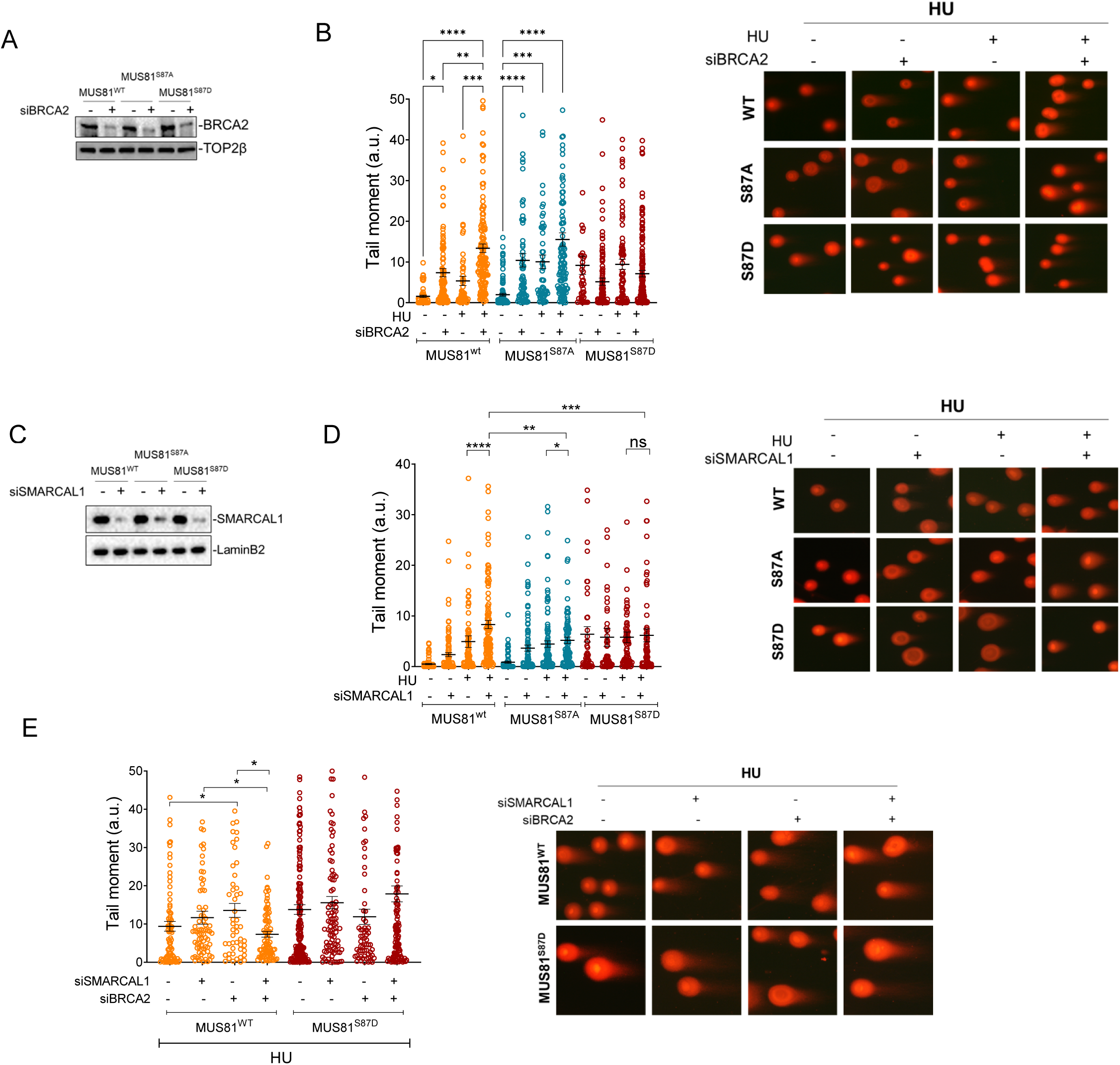
Deregulated function of MUS81 in S-phase exerted by the S87D mutation introduces DBSs independent of replication fork reversal. (A) Western blotting analysis of BRCA2 levels 72h after siRNA transfections in cells stably expressing the indicated MUS81 variant. TOP2β was used as loading control. (B) Analysis of DSBs by neutral Comet assays. Cells were transfected with BRCA2 siRNA and treated 72h later with 2m HU for 4h. Graph shows individual tail moment values from two different pooled replicates. Bars represent mean ± S.E. (C) Western blotting analysis of SMARCAL1 levels 48h after siRNA transfections in cells stably expressing the indicated MUS81 variant. LaminB1 was used as loading control. (D, E) Analysis of DSBs by neutral Comet assays. Cells were transfected with SMARCAL1 and/or BRCA2 siRNA and treated 72h later with 2m HU for 4h. Graph shows individual tail moment values from two different pooled replicates. Bars represent mean ± S.E. Statistical analyses were performed by Student’s t-test (*P< 0.5; **P< 0.1; ***P< 0.01; ****P< 0.001. Where not indicated, values are not significant).

Although expression of the inactive S87A MUS81 mitotic mutant increased the amount of DNA damage in BRCA2-proficient cells treated with HU, it did not affect the level of DSBs in cells depleted of BRCA2 and expressing wild-type MUS81 (Figure 4B).

Upon BRCA2 depletion, the wild-type MUS81 complex acts downstream fork reversal (31, 33). Since expression of S87D-MUS81 produced similar levels of DSBs irrespective of the BRCA2 status, we asked whether those DSBs were still dependent on fork reversal. Thus, we transiently depleted SMARCAL1 (Figure 4C), which is key for the reversal of the fork (35, 36), and evaluated if the formation of DSBs was affected similarly in cells expressing the wild-type MUS81 or the S87D mutant (Figure 4D). Depletion of SMARCAL1 in wild-type cells increased DSBs upon fork arrest, as previously observed (37). Interestingly, the presence of the mitotic S87A MUS81 mutant decreased the formation of DSBs in cells depleted of SMARCAL1, possibly suggesting that S87 phosphorylation is required to target replication intermediates accumulating when fork reversal is dampened. In contrast, depletion of SMARCAL1 did not affect DSBs in the S87D MUS81 mutant, suggesting that they accumulate independently of fork reversal. Consistent with this data, SMARCAL1 depletion failed to reduce DSBs in BRCA2-depleted cells expressing the S87D MUS81 mutant, while decreased their formation in the presence of the wild-type MUS81 (Figure 4E). To exclude that, in BRCA2-deficient cells expressing S87D-MUS81, DSBs could be induced by a different nuclease, we transfected these cells with a MUS81 siRNA that targeted the ectopic, shRNA-resistant, MUS81 or expressed the S87D-MUS81 bearing the nuclease-dead mutation (S87D+NUCD) and evaluated DSBs by neutral Comet assay (Supplementary Figure 3A-C). Transfection of MUS81 siRNA or introduction of the nuclease-dead mutation greatly reduced DSBs in the S87D MUS81 mutant confirming that the fork reversal-independent targeting of stalled forks is still dependent on MUS81 (Supplementary Figure 3B, C). Interestingly, however, reduction of DSBs in the S87D-MUS81 background was larger in cells depleted of MUS81 then in cells expressing the nuclease-dead mutation (Supplementary Figure 3B), possibly indicating that other nucleases might take over if the S87D MUS81 mutant is present.

Altogether, our results suggest that expression of the S87D MUS81 mutant, which mimics a deregulated function in S-phase of the mitotic MUS81/EME1 complex, counteracts fork degradation and exposure of parental ssDNA by cleaving stalled forks independently of fork reversal.

### Unrestrained activation of mitotic MUS81/EME1 complex prevents loading of fork reversal enzymes

We observe that the S87D MUS81 mutant induces DSBs also in the absence of SMARCAL1. Hence, we reasoned that recruitment of fork reversal factors would be reduced if breakage occurred before engagement of fork remodelling after replication arrest.

To this end, we analysed fork recruitment of SMARCAL1 and RAD51, both essential fork reversal factors, by *in situ* PLA assays and chromatin fractionation experiments.

Evaluation of the SMARCAL1 fork association by SIRF assay showed that HU treatment increased the presence of SMARCAL1 at the fork in wild-type cells, as expected (Figure 5A). SMARCAL1 was recruited at the stalled forks also in cells expressing the S87A MUS81 mutant, although the number of PLA spots were lower as compared with the wild-type (Figure 5A). In sharp contrast, fork recruitment of SMARCAL1 was severely reduced in cells expressing the S87D MUS81 mutant, either in untreated conditions or in response to HU (Figure 5A). Consistent with this result, the accumulation of detergent-resistant SMARCAL1 nuclear foci was greatly reduced in HU-treated cells expressing the S87D MUS81 mutant (Supplementary Figure 4A).

**Figure 5.**
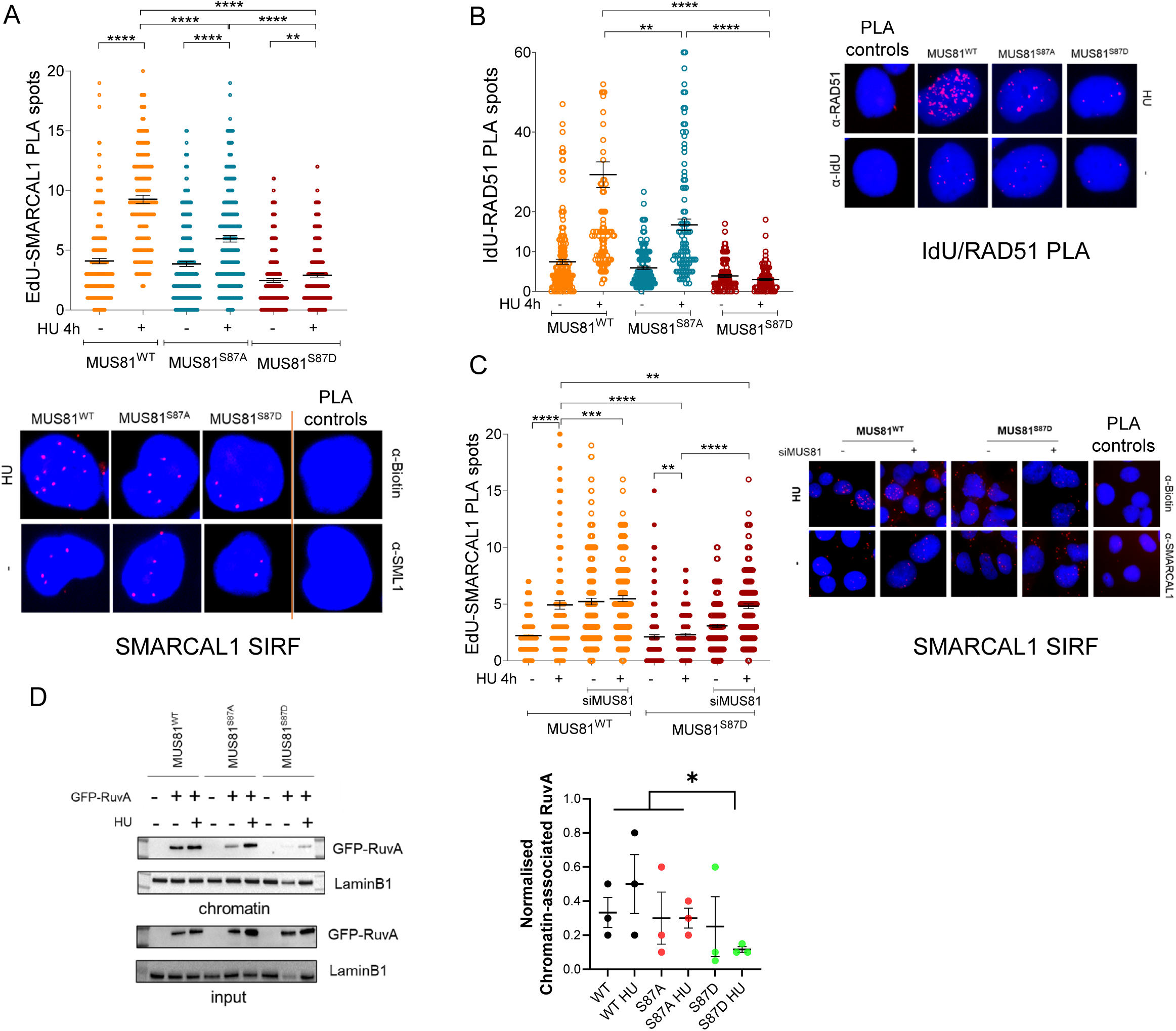
Deregulated function of the S87D MUS81 mutant affects correct fork recruitment of fork reversal factors. (A) SMARCAL1 interaction with active replication forks was detected by *in situ* PLA using anti-SMARCAL1 and anti-Biotin after labeling nascent DNA with EdU and Click-it reaction to conjugate Biotin to EdU. (B) RAD51 interaction with parental ssDNA was detected by *in situ* PLA using anti-RAD51 and anti-IdU after labeling parental DNA with IdU 20h before treatment. (C) SMARCAL1 interaction with active replication forks was detected as in (A) in cells transiently transfected with MUS81 siRNAs. The graphs show individual values of PLA spots from two pooled experiments. Representative images are shown. Bars represent mean ± S.E. (D) analysis of recruitment in chromatin of GFP-tagged RuvA. MRC5 shMUS81 cells stably expressing the indicated MUS81 variant were transfected with GFP-RuvA and chromatin fractions analysed 4h after treatment with 2mM HU by WB. The graph shows the normalized amount of GFP-RuvA in chromatin for two independent repeats. Statistical analyses were performed by ANOVA (**P< 0.1; ***P< 0.01; ****P< 0.001. If not indicated, values are not significant: p >0.5).

RAD51 functions upstream or downstream fork reversal (38) and analysis of fork recruitment by SIRF does not discriminate between those two roles. However, RAD51 loading at parental gaps at the stalled fork is reasonably involved in the promotion of fork reversal. Thus, to analyse the recruitment of RAD51 before fork reversal we used parental ssDNA-specific PLA (26). The presence of RAD51 at the parental ssDNA increased in response to HU in wild-type cells and, albeit at a slightly reduced level, also in the S87A MUS81 mutant (Figure 5B). However, the presence of RAD51 at the parental ssDNA was greatly reduced in the S87D MUS81 mutant. Similarly, formation of RAD51 foci was significantly decreased upon HU treatment in cells expressing S87D-MUS81 (Supplementary Figure 4B).

We hypothesised that recruitment of SMARCAL1 at fork in cells expressing S87D-MUS81 should be restored by blocking MUS81 if the impairment was dependent on aberrant cleavage before fork reversal. To test this prediction, we depleted MUS81 in cells expressing S87D-MUS81 and analysed fork recruitment of SMARCAL1 by SIRF (Figure 5C). In cells expressing wild-type MUS81, depletion of MUS81 increased recruitment of SMARCAL1 at fork in untreated cells but did not change the association in response to HU. Interestingly, depletion of MUS81 in cells expressing the S87D MUS81 mutant did not increase fork recruitment of SMARCAL1 in untreated cells but significantly enhanced the association after HU, restoring a wild-type behaviour.

To confirm our PLA and immunofluorescence data, we analysed localization in chromatin of SMARCAL1 and RAD51 by subcellular fractionation and Western blotting (Supplementary Figure 5). Treatment with HU for 4h increased the fraction of chromatin-associated SMARCAL1 and RAD51 in wild-type cells. In cells expressing the S87A MUS81 mutant, although the amount of SMARCAL1 and RAD51 in chromatin was lower compared to wild-type under unperturbed cell growth, both proteins were found enriched in chromatin after HU of about two fold. In striking contrast, the amount of SMARCAL1 and RAD51 associating with chromatin after HU was strongly reduced in cells expressing the deregulated S87D MUS81 mutant. Of note, the fraction of BRCA2 that associated with chromatin after HU was similar irrespective of the MUS81 status.

The formation of SMARCAL1-independent DSBs and the reduced recruitment of enzymes involved in fork reversal predicted that the number of reversed forks should be reduced in cells expressing S87D-MUS81. To test this hypothesis, we used recruitment in chromatin of the four-way junction-binding factor RuvA as a proxy for the presence of reversed forks (25). To this end, we ectopically expressed GFP-fused RuvA in cells expressing wild-type MUS81 or the two S87 mutants and induced replication fork stalling by a short HU treatment (Figure 5D). RuvA was chromatin-associated in all cell lines, consistent with the expected presence of a variable number of four-way junctions in the cells. After HU treatment, the normalised amount of RuvA in chromatin increased, although slightly, in cells expressing wild-type or S87A MUS81. In contrast, the normalised fraction of chromatin-associated RuvA was reduced in response to HU in cells expressing the S87D MUS81 mutant (Figure 5D).

It has been demonstrated that fork reversal slows down replication under treatments that perturb fork progression without inducing its complete arrest (39). Having demonstrate that the S87D-MUS81 mutant can target perturbed replication forks bypassing their reversal, we performed DNA fibre assays to evaluate if its expression could prevent the fork slowing associated with the fork reversal reaction. As shown in Supplementary Figure 6, expression of S87D-MUS81 did not prevent fork slowing under perturbed replication, possibly suggesting that introduction of DSBs is sufficient to slow down replication *per se*.

These results consistently indicate that S87D-MUS81, by escaping regulation in S-phase, targets unreversed replication forks after their stalling. Notwithstanding, S87D-MUS81 expression does not induce extensive cell death irrespective of the presence of BRCA2 and, consequently, HR.

This apparent inconsistency prompted us to analyse how DSBs were repaired in cells expressing S87D-MUS81. To this end, we performed neutral Comet assays in shMUS81 cells stably-complemented with MUS81 wild-type or S87D, in the presence or not of RNAi-mediated BRCA2 depletion (Figure 6A). Cells were exposed to HU to stimulate formation of DSBs and recovered in HU-free medium for 18h to promote their repair. Depletion of BRCA2 also served as control of the effect of an impaired HR-mediated repair while novobiocin was used to inhibit Polθ-dependent repair, which has been shown to be important for viability in the absence of BRCA2 and HR in general(40–42). To transiently-interfere with HR in untreated, BRCA2-proficient, cells, we used the B02 RAD51 inhibitor. As somehow expected, in untreated, wild-type cells, inhibition of RAD51 increased the amount of DSBs while it failed to stimulate DSBs in cells expressing S87D-MUS81 (Figure 6B). The small amount of DSBs seen in wild-type cells after HU treatment was repaired almost completely even in the presence of novobiocin, suggesting that multiple repair pathways can take over to deal with replication fork collapse induced or not by MUS81 (Figure 6B, C). Depletion of BRCA2 increased significantly the amount of DSBs induced by fork arrest but they are repaired, albeit more residual DNA damage was detected after recovery compared to CTRL-depleted cells (Figure 6B, C). In sharp contrast, cells expressing S87D-MUS81 accumulated DSBs after HU, which were insensitive to loss of HR (BRCA2-depletion) but were repaired almost completely after 18h (Figure 6B). Strikingly, however, S87D-MUS81-dependent DSBs were repaired predominantly by alt-EJ also in BRCA2-deficient cells (Figure 6B, C).

**Figure 6.**
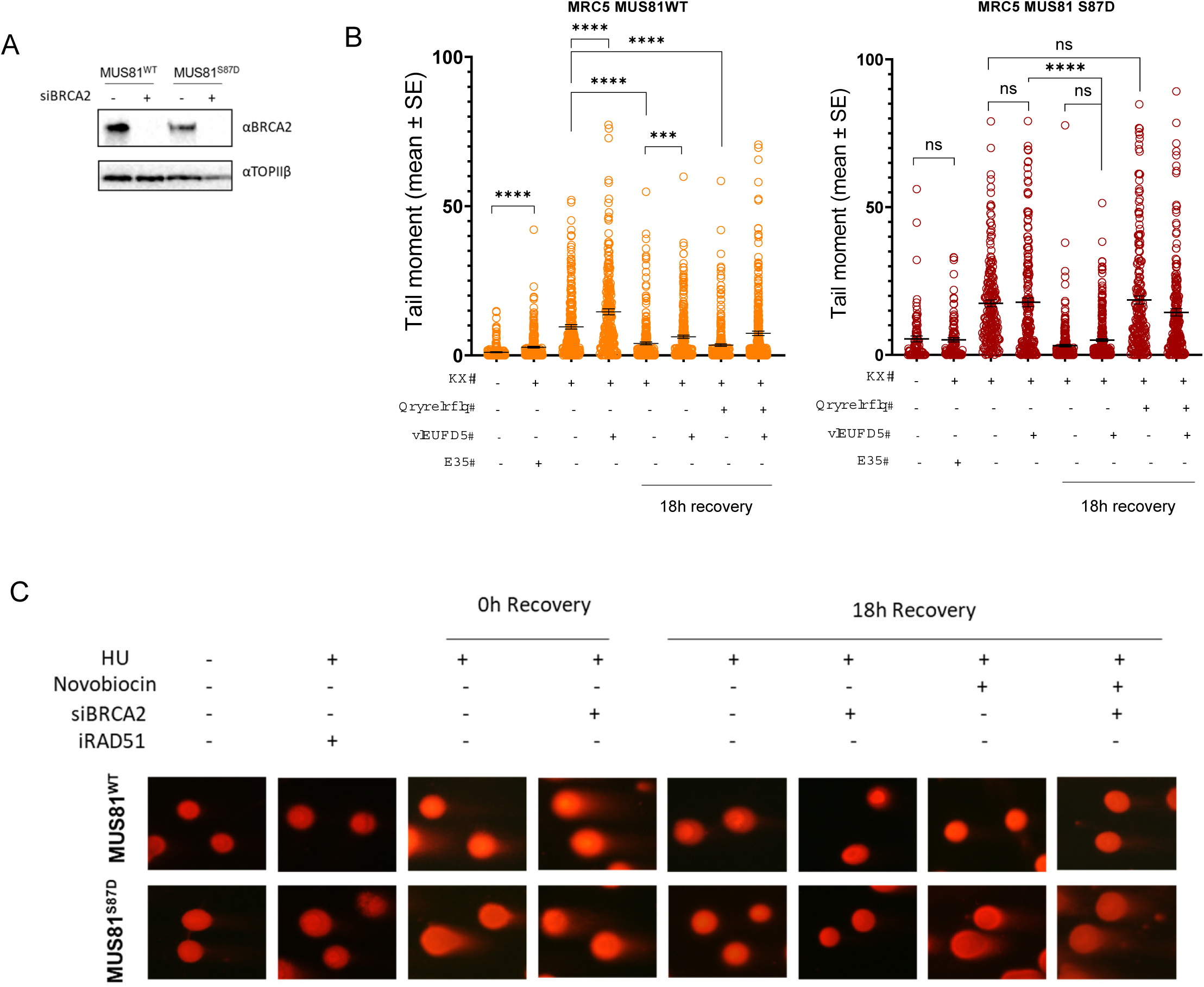
Deregulated function of MUS81 in S-phase exerted by the S87D mutation induces DBSs that are repaired by alt-EJ. (A) Western blotting analysis of BRCA2 levels 72h after siRNA transfections in cells stably expressing the indicated MUS81 variant. TOP2β was used as loading control. (B) Analysis of DSBs by neutral Comet assays. Cells were transfected with BRCA2 siRNA and treated 72h later with 2m HU for 4h (0h) and recovered for 18h in the presence of novobiocin. Graph shows individual tail moment values from two different pooled replicates. (C) Representative images are shown. Statistical analyses were performed by Student’s t-test (**P< 0.1; ***P< 0.01; ****P< 0.001. Where not indicated, values are not significant).

Altogether, these results indicate that the presence of the deregulated S87D MUS81 mutant greatly reduces the recruitment of crucial fork reversal factors at the stalled forks and also the chromatin association of RuvA, a proxy of four-way junctions, suggesting that unscheduled cleavage in S-phase by the mitotic-active form of the MUS81/EME1 complex takes place before fork remodelling. Moreover, our findings indicate that DSBs induced at unreversed forks by expressing S87D-MUS81 are repaired independently on HR even if BRCA2 is present, possibly explaining why S87D-MUS81 cells are viable upon BRCA2 depletion.

## DISCUSSION

Our findings show that disabling the mitotic function of the MUS81/EME1 complex is sufficient to induce enhanced cell death previously associated with MUS81 knockdown in BRCA2-depleted cells (21). Moreover, our data indicate that unscheduled activation of the MUS81/EME1 complex in S-phase conferred by the S87D-MUS81 mutant can largely revert Olaparib sensitivity in BRCA2-deficient cells.

These data may help explaining conflicting results on the effect of loss MUS81 on viability and chemosensitivity of BRCA2-deficient cells. Indeed, both MUS81 complexes, MUS81/EME1 and MUS81/EME2, are inactivated following downregulation of MUS81, but they might be differentially required in BRCA2-deficient cells. On one hand, the mitotic activity of the MUS81/EME1 complex might be involved in the resolution of late intermediates from BIR, which is over-active in the absence of BRCA2 (33, 43). On the other hand, MUS81/EME2 is thought to be involved in the processing of unprotected reversed forks (33). Our data using the S87A MUS81 mutant, which is a separation-of-function mutant disabling only the MUS81/EME1 function in mitosis (24), demonstrates that viability of BRCA2-deficient cells is primarily affected by inability to process persisting replication intermediates in G2/M irrespective of the normal ability in processing the deprotected, reversed, forks conferred by S87A MUS81 mutant, which is proven by the SMARCAL1-dependent formation of DSBs and expected fork degradation by MRE11.

Another debating feature of MUS81 concerns its relevance to chemosensitivity of BRCA2-deficient cells. Previous work demonstrated that abrogation of MUS81 function in BRCA2-deficient cells reduced sensitivity to PARPi treatment (20).

Strikingly, our data indicate that loss of the mitotic function of MUS81 does not affect sensitivity of BRCA2-deficient cells to Olaparib. Thus, response to Olaparib might be dependent on the ability of the MUS81/EME2 complex to process reversed forks in S-phase rather than on the processing of late DNA intermediates by the SLX4/MUS81/EME1 complex. Although deserving additional investigation, the apparent irrelevance of the mitotic MUS81 complex for the Olaparib sensitivity in BRCA2-deficient backgrounds might also suggest that the reported synergistic effect of WEE1i and Olaparib in breast tumour cells characterized by mutations in BRCA genes (44) does not derive from impaired activation of the SLX4/MUS81/EME1 complex, which is one of the WEE1-regulated targets (45).

Most strikingly, however, are our finding that Olaparib hypersensitivity of BRCA2-deficient cells can be largely reverted in the presence of the S87D MUS81 mutant. Indeed, the S87D MUS81 mutant mimics the activated status characteristic of the MUS81 mitotic complex leading to a normal functionality in late G2/M but is associated with a rogue functionality in S-phase (24). MUS81-dependent targeting of reversed and degraded forks is one of the determinants of Olaparib sensitivity in the absence of BRCA2 (20). Although apparently at odds with these observations, our data are supportive of previous models. Several of our findings are consistent with the hypothesis that deregulated MUS81/EME1-dependent cleavage occurs before fork reversal in BRCA2-deficient cells, and perhaps also in a the presence of BRCA2. Indeed, expression of S87D-MUS81 induces DSBs that are independent of SMARCAL1, one of the proteins essential for the reversal of stalled forks (36), and consistently impairs the replication fork arrest-dependent recruitment of SMARCAL1 and RAD51. Moreover, expression of S87D-MUS81 decreases accumulation in chromatin of the 4-way junction binder RuvA, a proxy for the presence of reversed forks, suggesting that less reversed forks are formed upon replication fork stalling. Consistently, the expression of S87D-MUS81 counteracts fork degradation by MRE11 in BRCA2-deficient cells, an event that occurs downstream fork reversal and that is abrogated whenever reversion of the fork is blocked (31, 33, 34). Interestingly, inactivation of the nuclease activity in the S87D-MUS81 still prevents MRE11-dependent fork degradation in the absence of BRCA2. This might be at odds with the hypothesised role of de-regulated cleavage induced by the S87D mutation but can be explained with take-over by other nucleases capable of targeting unreversed forks, as suggested by residual DSBs observed in this double mutant. Indeed, under pathological conditions, depletion of MUS81 does not prevent DSBs completely because SLX4 takes over (15), and SLX4/SLX1 can target unreversed forks when SMARCAL1 is depleted (37, 46). Since the S87D mutation in MUS81 promotes association with SLX4 in mitosis (24), it tempting to speculate that even a catalytically-dead S87D-MUS81 might retain or attract SLX4 at fork promoting its take-over. This is an interesting point to address in future studies. Alternatively, requirement of a functional MUS81 for the processing of 5’-flaps at degraded forks and for the reversal-degradation cycle that is behind the appearance of shorten replication tracts in the DNA fiber assay might mask rescue of fork degradation in cells expressing the nuclease-dead S87D MUS81 mutant (47).

Thus, resistance to Olaparib induced by deregulated MUS81/EME1-dependent cleavage in S-phase would derive from introduction of DSBs at stalled replication forks before they can be processed by fork reversal enzymes (Fig. 7). This mechanism casts the possibility that unreversed, stalled, forks might become substrate for MUS81/EME1 in a complex with SLX4. Consistent with this possibility, SLX4-dependent DSBs are formed when SMARCAL1 is not functional (37, 46) and we found that expression of S87A-MUS81, which confers reduced ability to interact with SLX4 (24), decreases the formation of DSBs in SMARCAL1-depleted cells. In a BRCA2 wild-type background, those DSBs can be handled and repaired by HR, as indicated by a proficient BRCA2 focus-forming activity in cells expressing the S87D-MUS81. However, in the absence of BRCA2, they can be likely repaired by other pathways given that S87D-MUS81 expression in BRCA2-deficient cells does not affect viability. Consistent with activation of alternative repair pathways, DSBs induced by the S87D MUS81 mutant are repaired preferentially by alt-EJ and not by multiple repair pathways, as it occurs in wild-type cells (Fig. 7). Engagement of an HR-independent repair pathway of the S87D-MUS81-dependent DSBs is also consistent with the observation that BRCA2 foci were not significantly higher than in the S87A-MUS81 mutant despite the presence of about 10 times more DSBs.

**Figure 7.**
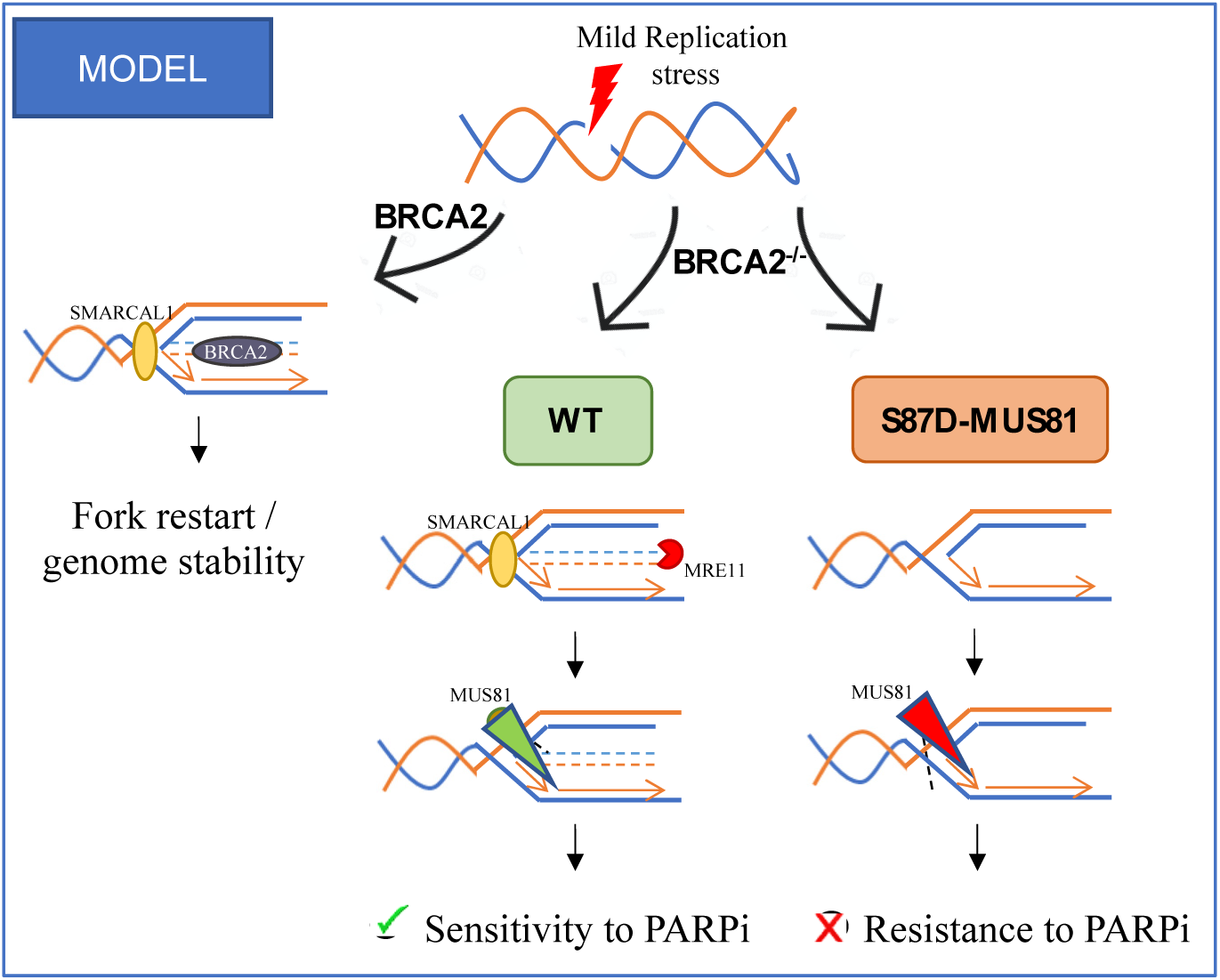
Proposed model of the S87D MUS81 mutant action at perturbed replication forks in normal or BRCA2-deficient cells and implications for PARPi resistance. After replication fork stalling, BRCA2 acts to protect reversed forks from MRE11-mediated degradation, ensuring correct fork restart and genome stability. In the absence of BRCA2, reversed forks are deprotected and undergo degradation followed by MUS81-dependent DSBs. This, together with accumulation of gaps, contributes to PARPi sensitivity. In the presence of unscheduled function of MUS81/EME1 conferred by S87D-MUS81, perturbed replication forks undergo breakage before fork remodelling, and this confers resistance to PARPi by stimulating a Polθ-dependent DNA repair pathway.

Thus, the ability of S87D-MUS81 to bypass fork degradation and HR-dependent repair in BRCA2-deficient cells is sufficient to overcome Olaparib sensitivity. Interestingly, a quick conversion of perturbed replication forks into DSBs might also counteract accumulation of parental ssDNA gaps, a phenotype that has been recently correlated to PARPi sensitivity in BRCA-deficient cells (8). Indeed, expression of S87D-MUS81 greatly reduces the presence of parental ssDNA in response to replication fork arrest in the absence of BRAC2 and this reduction might also derive from less template gaps, in addition to an impaired degradation of the regressed arm of reversed forks.

MUS81-dependent cleavage at perturbed replication forks before their reversal has been demonstrated recently in response to transcription/replication conflicts (48). In that case, breaks introduced by MUS81 are repaired using a Lig4/XRCC4-dependent pathway but out of the NHEJ (48). We observe that S87D-MUS81-dependent DSBs occurring independently of fork reversal are repaired almost entirely by a Polθ-dependent mechanism, likely alt-EJ. Thus, deregulated action of MUS81 or its role in response to transcription/replication conflicts defines two distinct mechanisms. Our results may also have strong translational implications for cancer therapy. Indeed, CK2 has been found overexpressed in many human tumours (49). Thus, although cancer-related mutations leading to deregulated function of the SLX4/MUS81/EME1 complex in S-phase have not been reported yet, it is tempting to speculate that in CK2-overexpressing cancer cells also the biological function of MUS81-EME1 may be elevated contributing to chemoresistance and, possibly, increased genome instability and aggressiveness. Also PLK1 might contribute to activation of MUS81/EME1, and PLK1 is often deregulated in cancers (50, 51). It would be worth investigating if Olaparib-resistance is more frequent in cancers with increased levels of SLX4/MUS81/EME1 regulatory kinases.

Altogether, our study defines the specific role of MUS81 implicated in the synthetic-sick genetic relationship with loss of BRCA2 and provides the first mechanistic insight into chemoresistance associated with deregulated activity of MUS81/EME1 in BRCA2-deficient cells.

## ACKNOWLEDGMENTS

We thank Profs. Daniela Barilà and Marco Barchi for helpful discussion and advise. This work was supported by Italian Ministry of Health “Ricerca Finalizzata” to P.P. (grant n. RF-2016-02362022) and by investigator grants from Associazione Italiana per la Ricerca sul Cancro (AIRC) to P.P. (IG n. 21428) and to A.F. (IG n. 19971).

## AUTHORS CONTRIBUTION

F.B. performed experiments to analyse Olaparib sensitivity and experiments to assess the mechanism of chemoresistance. E.M. performed experiments to evaluate RuvA chromatin loading, DNA fibers and DNA degradation experiments. G.M.P performed experiments to evaluate mitotic defects in BRCA2-depleted cells and the initial assessment of viability in MUS81 mutants. C.F. performed the viability analyses upon XRCC4 depletion and DSBs repair assays. A.N. performed experiments with nuclease-dead MUS81. All authors contributed to design experiments and analyze data. P.P and A.F. designed experiments, analysed data and supervised the project. F.B. contributed to write results. P.P. and A.F. wrote the paper. All authors contributed to revise the paper.

## CONFLICT OF INTEREST

The authors declare to do not have any conflict of interest

## Notes

### Competing Interest Statement

The authors have declared no competing interest.

### Summary of Updates

New experiments to analyse repair of S87D-MUS81-induced DSBs and additional experiments to include the phenotype of S87D-MUS81 with additional mutations to impair nuclease function.

