## Supplementary Figures and Legends for "Phosphorylation Status Of MUS81 Is A Modifier Of Olaparib Sensitivity In BRCA2-Deficient Cells"

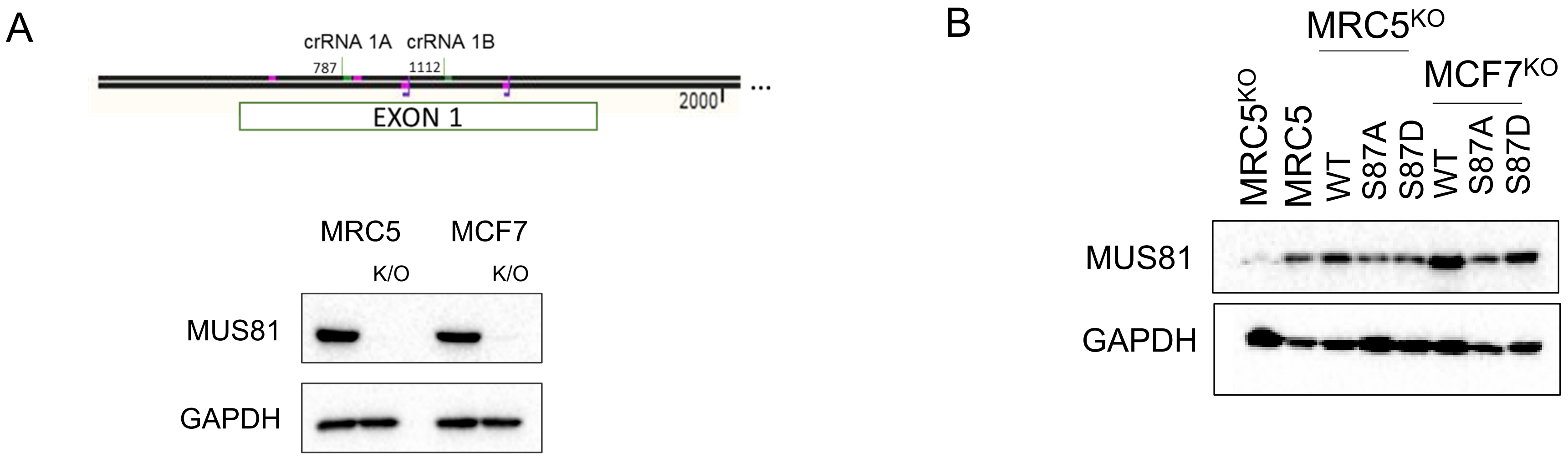


**Supplementary Figure 1. Generation of MUS81 KO cell lines.** (A) Scheme depicting the position of the two Cas9 targeted site in the MUS81 exon 1. Western blotting analysis shows the level of MUS81 protein after knockout from two isolated clones. (B) Western blotting shows the levels of MUS81 protein after transfection with the indicated MUS81 constructs in the indicated MUS81 KO clones. GAPDH was used as loading control.

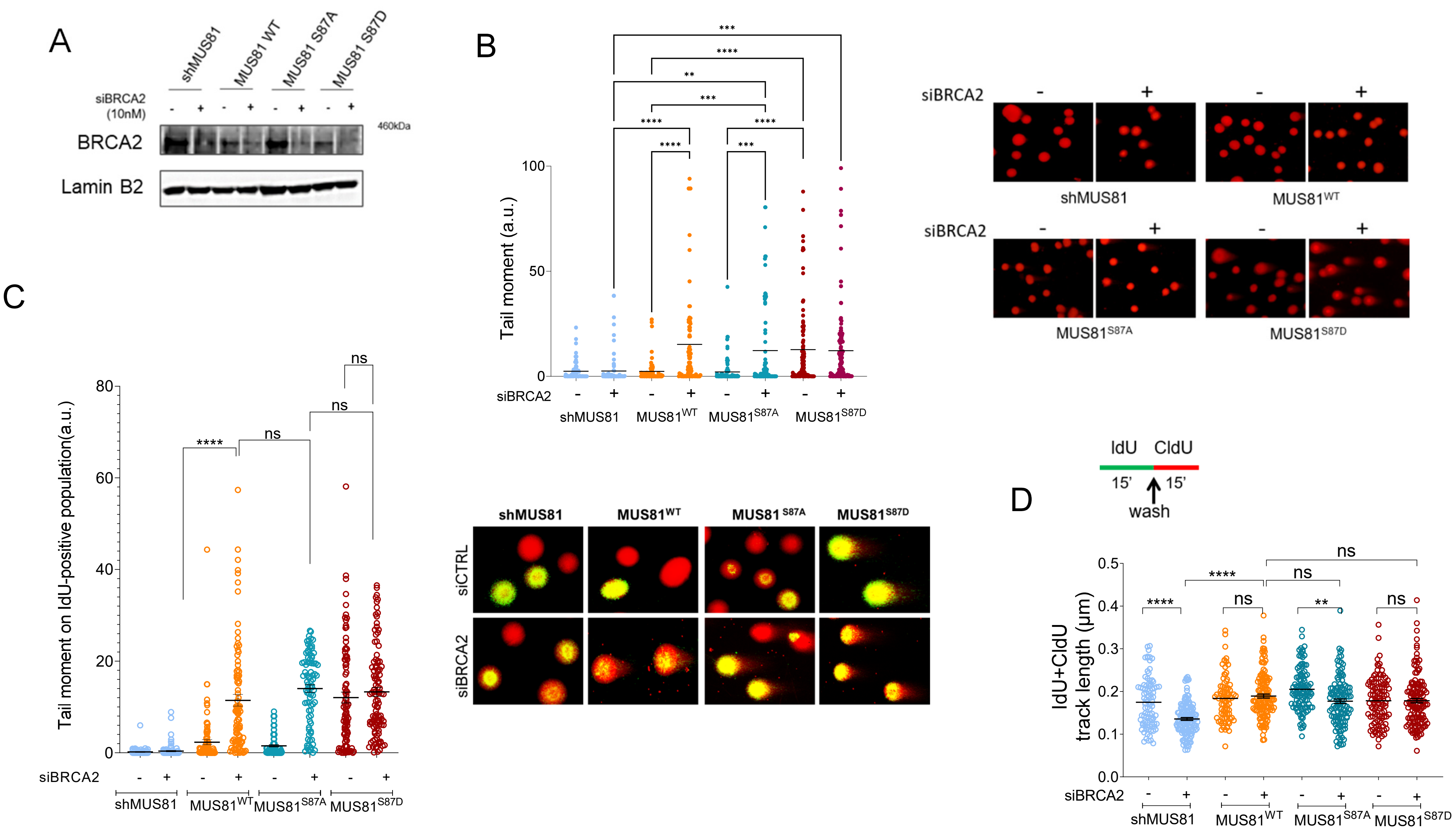


**Supplementary Figure 2. Formation of DSBs in BRCA2-deficient cells does not depend on the mitotic function of MUS81.** (A) Western blotting analysis shows level of BRCA2 protein after transfection with siRNA. Lamin B1 was used as loading. (B) Analysis of DSB accumulation by neutral Comet assay. Dot plot shows data presented as individual tail moment from three independent pooled experiments. Representative images are shown. (C) Formation of DSBs in S-phase after BRCA2-depletion. Cells were transfected as indicated. After 48h cells were labeled with IdU and DSBs were evaluated by neutral Comet assay followed by IdU immunofluorescence. Tail moment was quantified on IdU positive cells. Representative images of Comets are presented in the panel (D) Experimental scheme of dual labelling replication assay for DNA fibres. Red tract: CldU; green tract: IdU. Dot plot showing the IdU+CldU tract length (µm) of ongoing forks in single DNA fibres. The length of the tract was measured in at least 100 well isolated DNA fibres from three independent experiments. Mean values are represented as horizontal black lines ± S.E. (ns, not significant; *P< 0.1; **P< 0.01; ***P< 0.001 ****P< 0.0001; Mann–Whitney test).

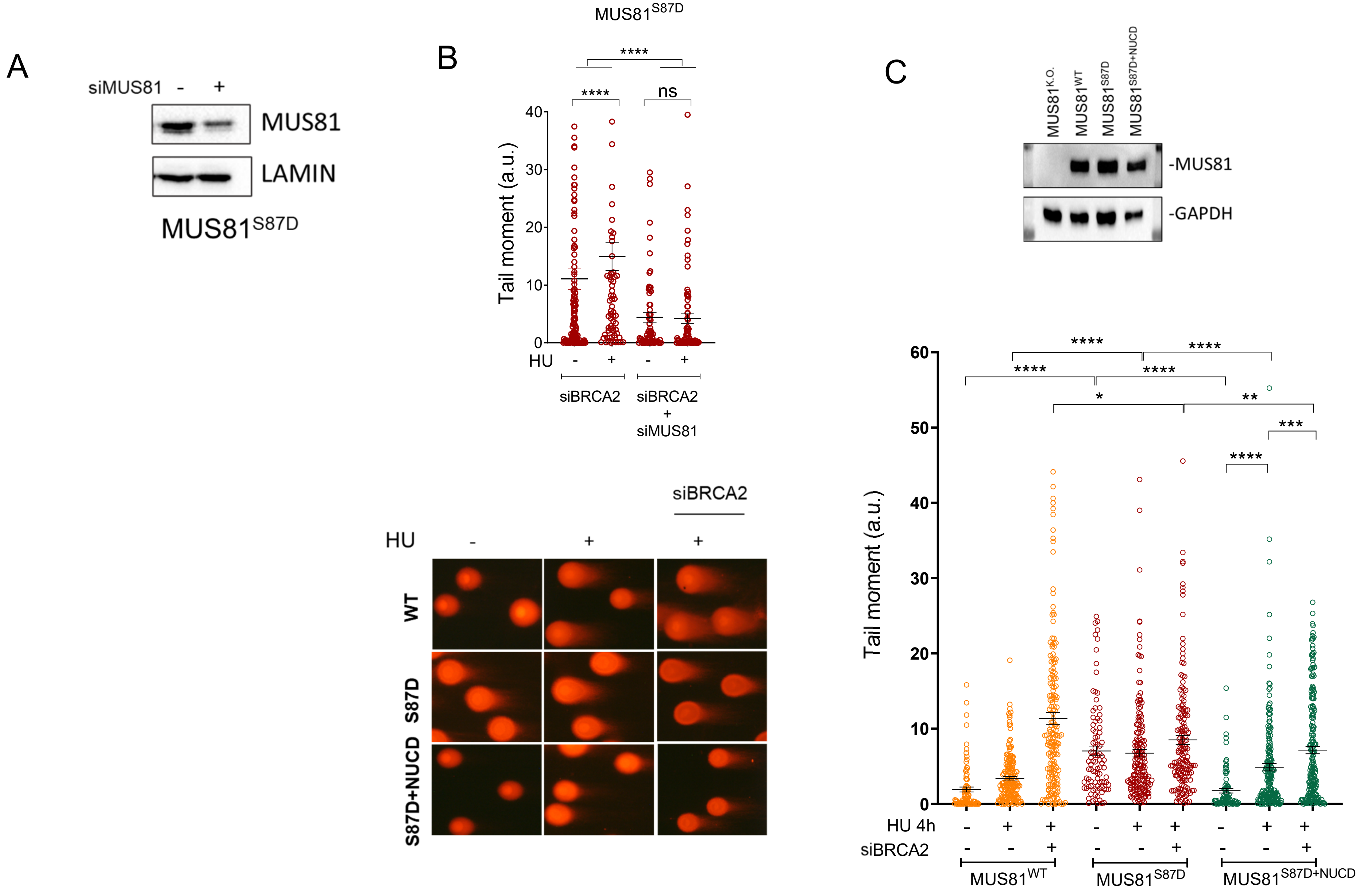


**Supplementary Figure 3. DSBs in BRCA2-deficient cells expressing the S87D MUS81 mutant are still dependent on MUS81** (A) Western blotting analysis shows level of MUS81 protein after transfection with siRNA. Lamin B1 was used as loading. (B)(C) Analysis of DSB accumulation by neutral Comet assay. Dot plot shows data presented as individual tail moment from two independent pooled experiments. Mean values are represented as horizontal black lines ± S.E. S87D+NUCD = S87D mutant combined with the two nuclease-inactivating mutations (D338A/D339A) (ns, not significant; ****P< 0.0001; Student's t-test).

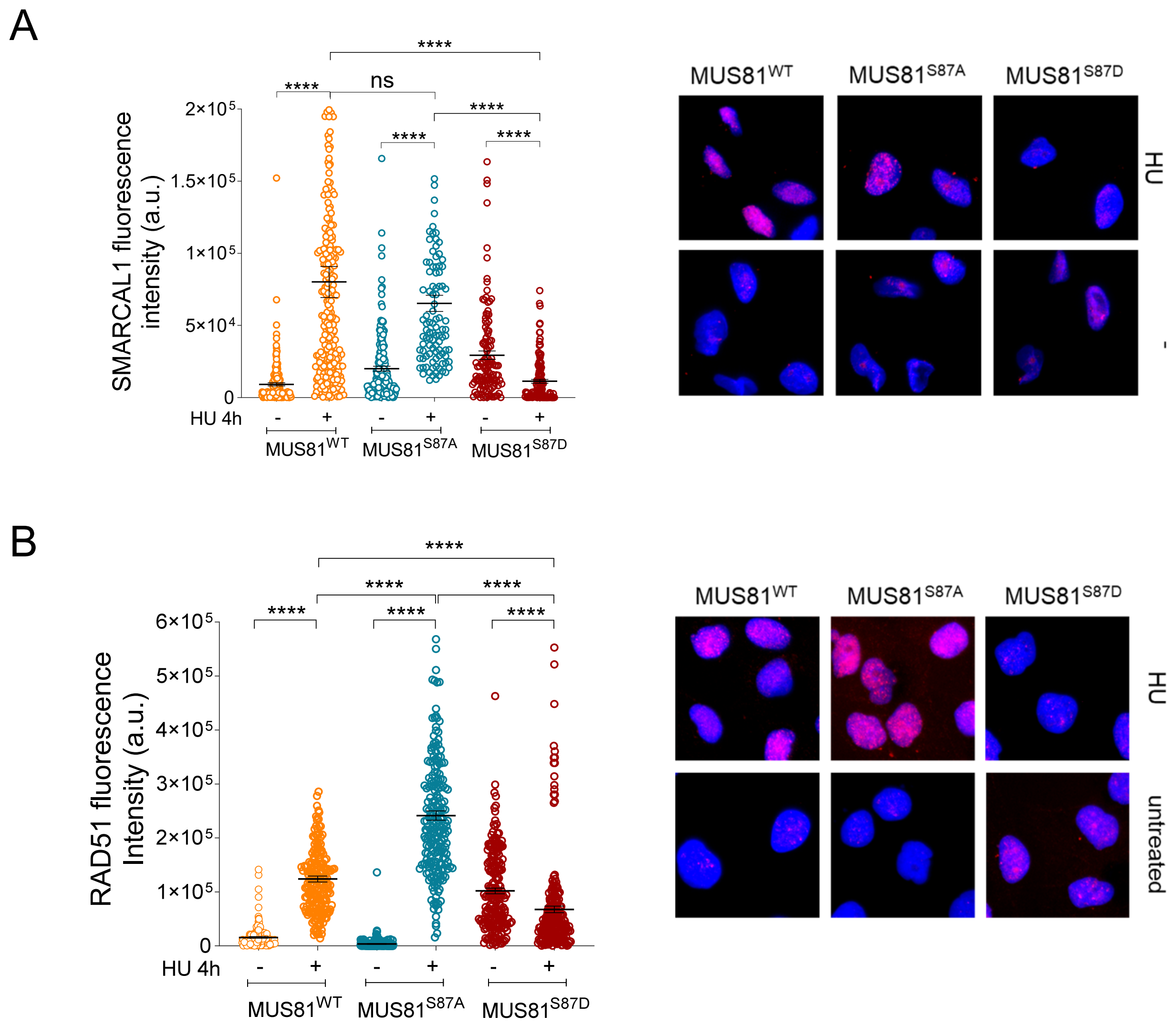


**Supplementary Figure 4. Analysis of SMARCAL1 and RAD51 foci formation in cells expressing the S87 MUS81 mutants.** Anti-SMARCAL1 (A) or anti-RAD51 immunofluorescence staining (B) was performed in MRC5 shMUS81 cells stably complemented with the indicated MUS81 variants. Graphs show quantification of the fluorescence intensity from at least 200 nuclei (n=2); horizontal black lines represent the mean ± S.E. Representative images were shown. Statistical analysis was performed by Student's t-test. (ns: not significant; ***P<0.001).

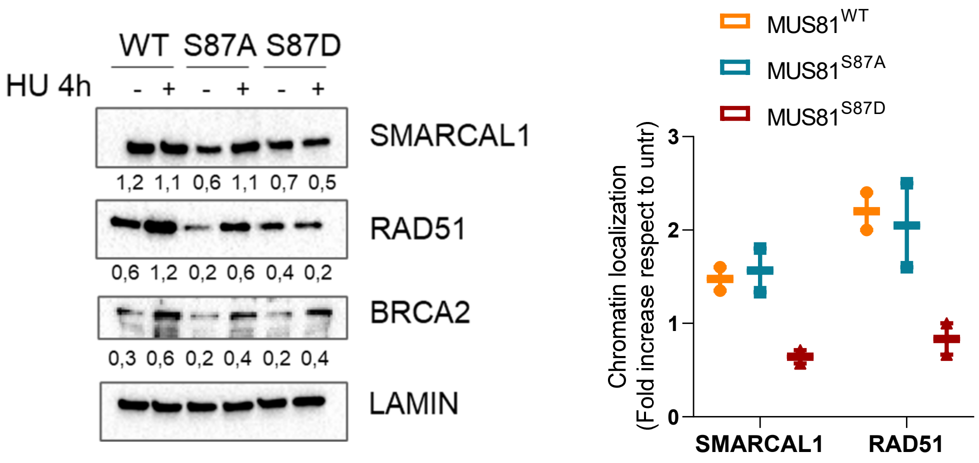


**Supplementary Figure 5. Analysis of SMARCAL1 and RAD51 chromatin recruitment in cells expressing the S87 MUS81 mutants.** Chromatin fractionation assay was performed in MRC5 shMUS81 cells stably complemented with the indicated MUS81 variant and treated with 2mM HU. The membrane was probed with the indicated antibodies. The amount the chromatin-bound proteins are normalized against LaminB1 and reported in the graph as fold increase over the untreated.

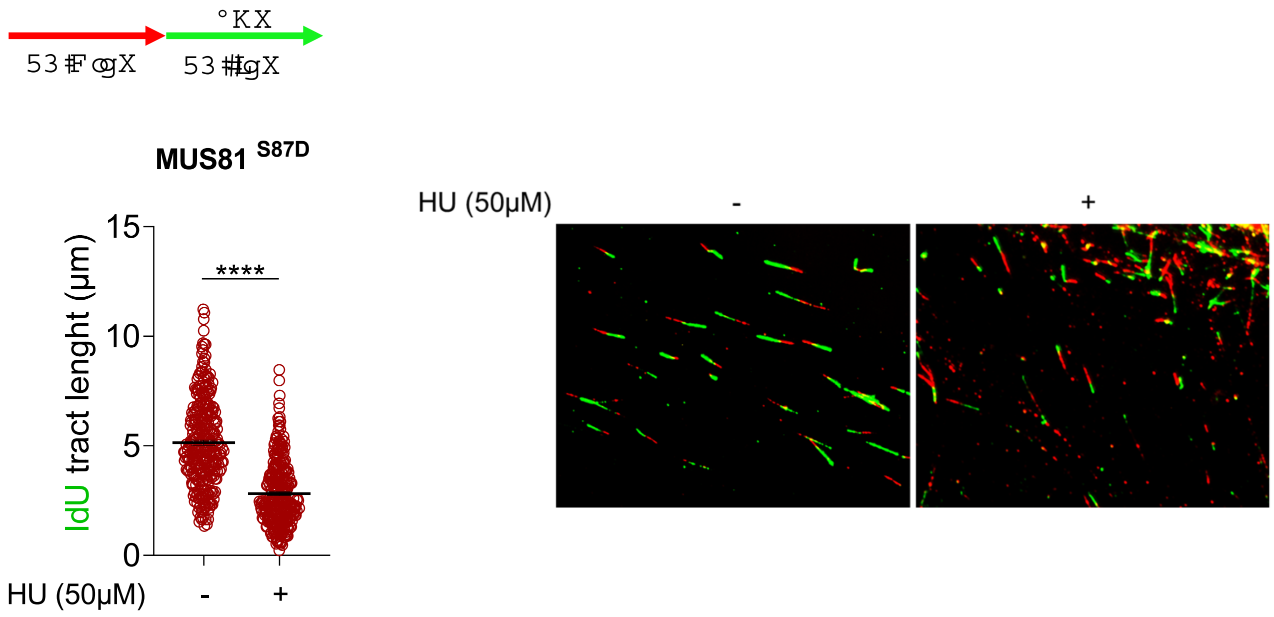


**Supplementary Figure 6. Analysis of fork progression in cells expressing the S87D MUS81 mutation. SMARCAL1 and RAD51 chromatin recruitment in cells expressing the S87 MUS81 mutants.** shMUS81 MRC5 cells stably complemented with the S87D MUS81 mutant were treated as indicated in the scheme on top. The graph shows length of IdU tract in presence or not of 50µM hydroxyurea treatment. Representative images of DNA fibers fields are presented. Values from at least 150 fibers are plotted and black lines represent means ± SE (****P < 0.001; Mann–Whitney test).
